# Can you affect me? The influence of vitality forms on action perception and motor response

**DOI:** 10.1101/2021.05.04.442561

**Authors:** G. Lombardi, J. Zenzeri, G. Belgiovine, F. Vannucci, F. Rea, A. Sciutti, G. Di Cesare

## Abstract

During the interaction with others, action, speech, and touches can communicate positive, neutral, or negative attitudes. Offering an apple can be gentle or rude, a caress can be kind or rushed. These subtle aspects of social communication have been named *vitality forms* by Daniel Stern. Although they characterize all human interactions, to date it is not clear whether vitality forms expressed by an agent affect the action perception and the motor response of the receiver. To this purpose, we carried out a psychophysics study aiming to investigate how perceiving different vitality forms can influence cognitive and motor tasks performed by participants. In particular, participants were stimulated with requests made through a physical contact or vocally and conveying rude or gentle vitality forms, and then they were asked to estimate the end of a passing action observed in a monitor (action estimation task) or to perform an action in front of it (action execution task) with the intention to pass an object to the other person presented in the video. Results showed that the perception of different vitality forms influences both the estimated duration of the action and the motor response of participants, suggesting how these forms of communication can positively or negatively affect our behavior.

## Introduction

Social interactions are characterized by the capacity to communicate our intentions and affective states, and to evaluate those of others. This behavioral exchange is based on actions and speech dynamics of the interactants which have been defined by Daniel Stern with the term “vitality forms”.^1^ Vitality forms play a bidirectional role in human-human interactions: the perception of vitality forms allows the receiver to understand the affective state of the agent while the expression of vitality forms allows the agent to communicate his mood/attitude.^2^ For example, a simple action (e.g., passing an object) can be performed gently or rudely, according to the positive or negative attitude of the agent.

In recent years, fMRI studies have investigated the neural substrates of these forms of communication, showing that the observation, imagination, and execution of actions conveying vitality forms induce the activity of the dorso-central insula.^2-6^ Moreover, the same authors have demonstrated that the perception of speech and action sounds (i.e. stir the coffee, knock the door, etc.) and expression of words conveying vitality forms activate the same insular sector.^5-8^ Several authors^9-11^ suggest that action understanding is reached by an internal motor simulation in which the observed action is remapped on the observer’s motor schema allowing them to internally replicate the action goal. It is plausible that during social interactions, the physical and the affective component of actions (vitality forms) may be reproduced internally, showing a contagion effect on the subsequent motor response.^12-15^ This hypothesis is corroborated by a kinematic study carried out by Di Cesare and collegues,^16^ showing that a gentle and rude request presented as visual actions (visual modality) or spoken action verbs (auditory modality) or both (mixed modality) influenced the motor response of participants. In particular, vitality forms modulated spatial and temporal parameters of actions, highlighting a larger trajectory and higher velocity in response to rude requests, compared to the gentle ones.

In everyday life, besides actions and speech, individuals also interact using physical touch. A typical example is a common greeting gesture such as handshaking. Indeed, the handshake conveys information about the affective state of the individual via touch and proprioception.^17^ In this view, an interesting question is to understand, besides action and speech, whether a physical touch may also influence the receiver. For this purpose, in the present study we investigate: 1) how a rude or gentle request expressed through physical contact or vocally by an actor can influence the perception of actions; 2) how the same requests can modulate the kinematic parameters of a subsequent action. Specifically, participants were required to perform two different tasks. In the first task (Action Estimation Task), they were presented with video clips showing the initial part of a passing action performed with different vitality forms (rude or gentle) and were asked to estimate the time of its completion by pressing a button. Before the videos’ presentation participants received two different types of stimulation: a request mediated by physical contact, during which the robotic manipulandum reproduced a rude or gentle movement on their right arm, or a vocal request (“give me”) pronounced with a rude or gentle voice by two actors (a male and a female). In the second task (Action Execution Task), participants received a request mediated by physical contact or vocally conveying rude or gentle vitality form and then were required to move actively the handle of the manipulandum with the aim to pass the object to another person showed on the monitor. We hypothesized: 1) an effect of the perception of different vitality forms on the estimation of action duration; 2) an effect of the perception of different vitality forms on the execution of actions for both the vocal stimuli (as shown in a previous kinematic study^16^) and the physical contact.

## 2. Materials and Methods

### 2.1 Participants

The experiment was carried out on 18 healthy right-handed volunteers (twelve females and six males; mean age = 24.1 years; SD = 2.7 years). All participants were native Italian speakers and had normal or corrected-to-normal vision and normal hearing. None reported neurological or cognitive disorders.

### 2.2 Tasks and Experimental Paradigm

Participants sat in a comfortable chair in front of a monitor, holding the handle of the manipulandum “Braccio di ferro”^18^ with their right hand and wearing a pair of headphones (Figure 1 B1). The monitor was set to a spatial resolution of 1920 × 1080 pixels. The participants were required to perform two main tasks: an *Action Estimation Task* and an *Action Execution Task*. In the *Action Estimation Task* participants observed videos in which the right hand of an actor passed an object (a ball, a bottle, a cup or a packet of crackers) to another person represented on the other side of a table (Figure 1 A), with either a gentle or a rude vitality form.

**Figure 1.**
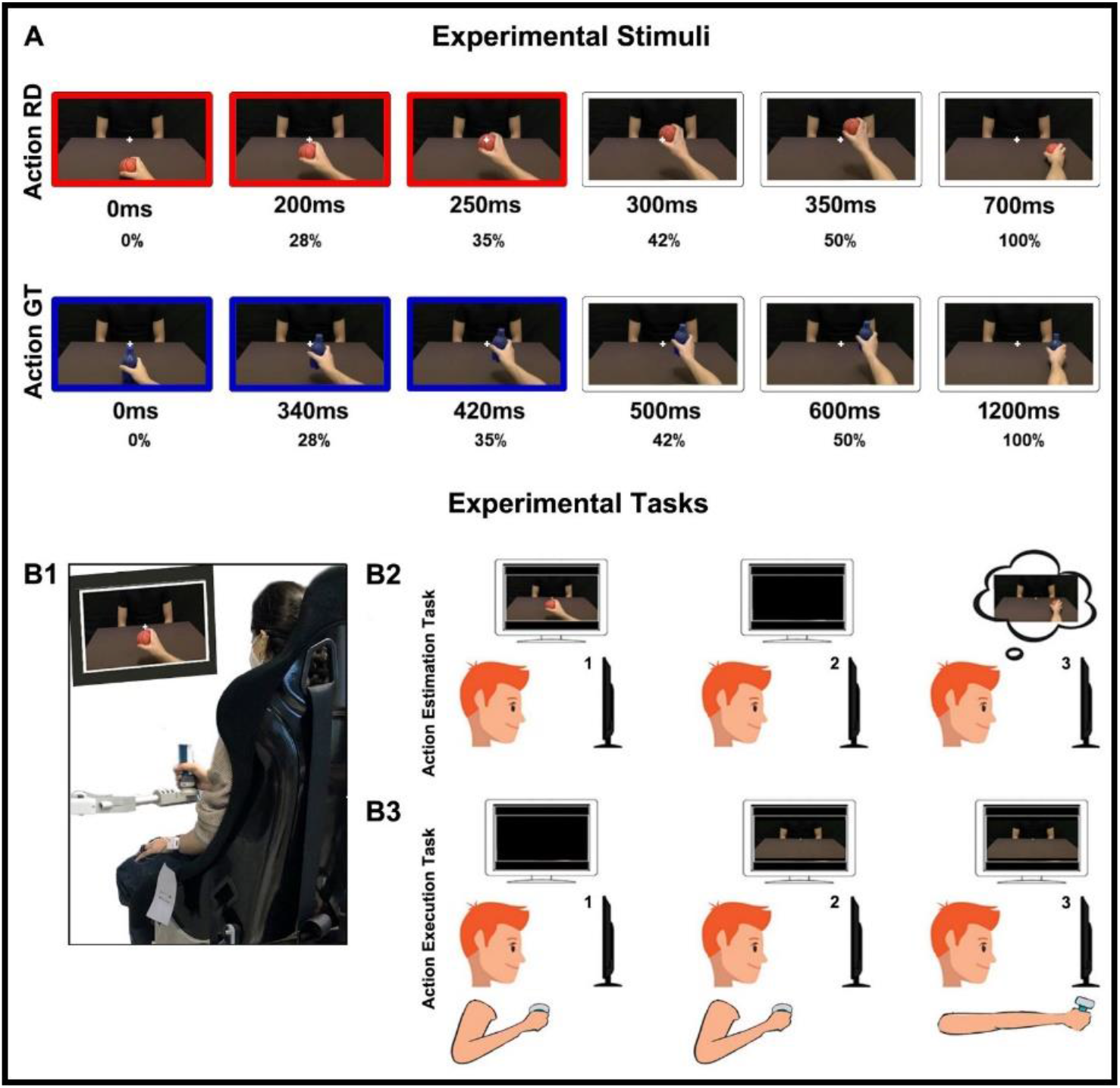
Visual stimuli (A): the total duration was respectively 700ms for rude actions and 1200ms for gentle actions. The stimuli consisted in presenting the 35% of the entire duration: 250ms for rude actions (red color) and 420ms for gentle actions (blue color). Example of a participant during the experiment (B1). *Action Estimation Task* (B2): 1. participants observed the initial part of a passing action, 2. the action was obscured, 3. they continued the action mentally estimating the time of its conclusion. *Action Execution Task* (B3): 1. starting position, 2. a static image of an actor appeared, 3. participants performed the passage moving the handle.

This egocentric perspective allowed participants to get involved in the action. More specifically, video stimuli consisted of showing only an initial part (35%) of the entire passing action, corresponding to 250ms for rude actions (Figure 1 A, red color) and to 420ms for gentle actions (Figure 1 A, blue color). After these durations, actions were obscured and participants were asked to continue them mentally, indicating the time of their conclusion by pressing a button located on the handle of the manipulandum (Figure 1 B2). In the *Action Execution Task* participants were instead presented with a static image of the same actors and were required to move actively the handle in front of them with the intention to pass an object. (Figure 1 B3). The experiment was composed of five runs (Figure 2). In the Baseline run participants simply estimated the duration of actions. In the Estimation Physical Request run a physical request preceded the action estimation task: first, the manipulandum moved the participants’ arm gently or rudely and subsequently, they observed the initial part of the action and estimated its end. In the Action Physical Request run after receiving the physical request executed gently or rudely by the manipulandum, participants performed the action execution task, moving actively the handle towards the actor/actress. In the remaining runs requests were vocal. In particular, in the Estimation Vocal Request run participants listened to a voice pronouncing “give me” in a gentle or rude way and then they executed the action estimation task. Finally, in the Action Vocal Request run after receiving the vocal request expressed gently or rudely, participants performed the action execution task. With the purpose of avoiding a bias due to the runs order effect, the presentation order of experimental runs was balanced across participants. Nine participants started with the runs presenting a vocal request followed by the runs presenting a physical request. Nine participants started with the runs presenting a physical request followed by the runs presenting a vocal request. The stimuli presented randomly in the Baseline run were 24: 12 rude actions and 12 gentle actions, presented at the beginning of the experiment. In each of the two “Estimation” runs a random presentation of congruent and incongruent conditions was created to evaluate the influence of vitality forms characterizing the request on the action estimation task. This means that the requests (physical or vocal) and the following action stimuli could share the same vitality form (*rude* request – *rude* action or *gentle* request – *gentle* action) or could have opposite vitality form (*rude* request – *gentle* action or *gentle* request – *rude* action). In particular, each of these runs presented 48 stimuli consisting of: 12 rude requests followed by rude actions (congruent condition), 12 rude requests followed by gentle actions (incongruent condition), 12 gentle requests followed by gentle actions (congruent condition), 12 gentle requests followed by rude actions (incongruent condition).

**Figure 2.**
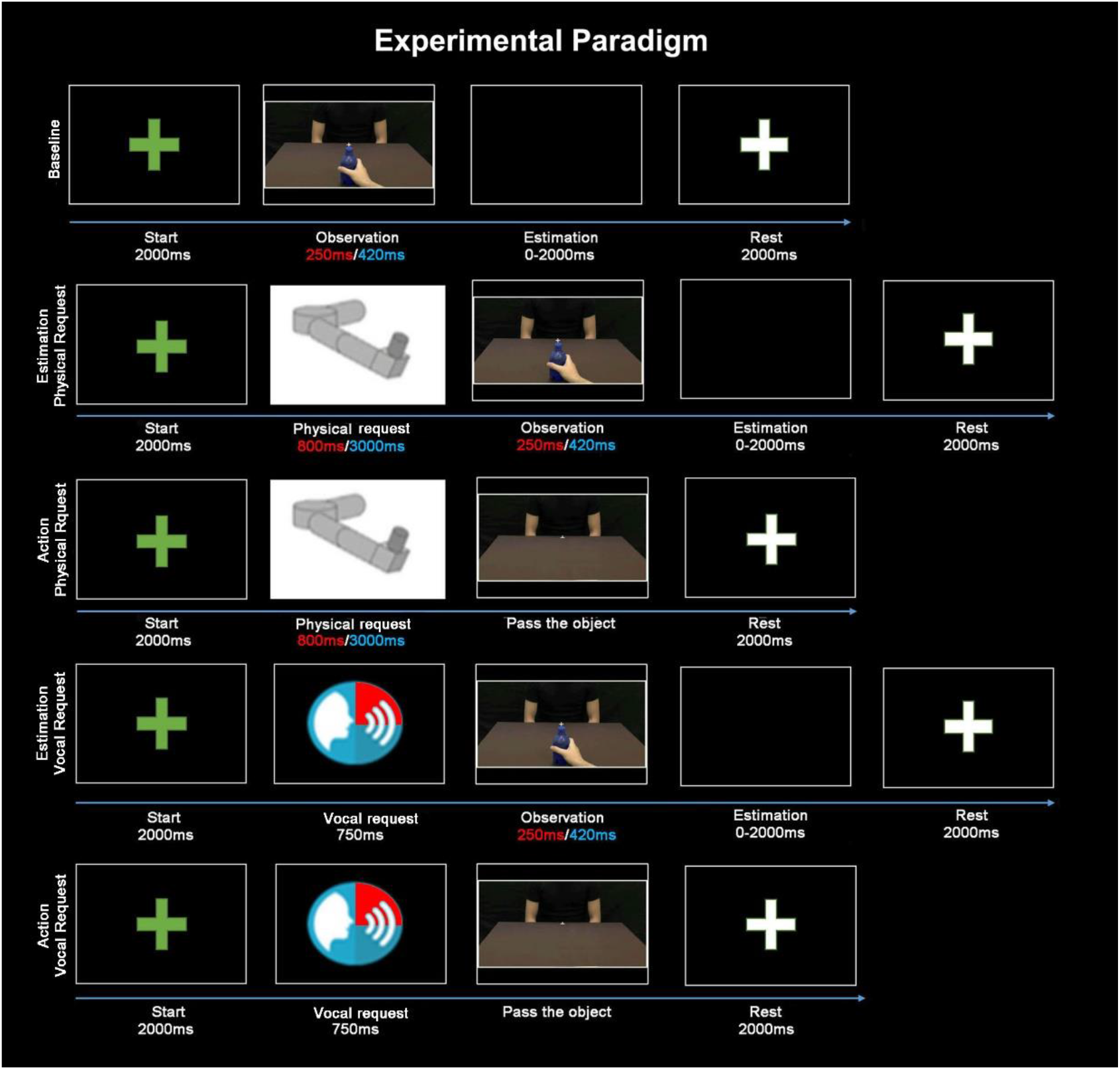
For each run, the green fixation cross (Start, 2000ms) indicated the beginning of a new trial. In the Baseline run and in the two Estimation runs (2^nd^ and 4^th^ rows) participants observed the initial part of the action (Observation), then the monitor turned black (Estimation, 0-2000ms), and when they pressed the button on the handle to estimate the action conclusion a white fixation cross appeared (Rest, 2000ms). In the two Action runs (3^rd^ and 5^th^ rows), when a static image of an actor/actress appeared (Pass the object), participants moved actively the handle in front of them. At the end of the passage, the manipulandum moved passively the arm at the starting position and the white fixation cross appeared (Rest, 2000ms). The panel with the manipulandum icon (Physical Request) indicated that a physical request preceded the subsequent task. The panel with the audio icon indicated that a vocal request preceded the subsequent task. Red color corresponds to rude vitality forms while blue color corresponds to gentle vitality forms.

Finally, each of the two “Action” runs presented in a random order 24 stimuli: 12 rude requests and 12 gentle requests. For each run, a period of rest of two seconds was inserted between trials and it was marked by a white fixation cross on a black background to keep attention on the screen. Before the beginning of a new trial the white cross turned green. PsychoPy v3.0 software was used to present video stimuli and to record participants’ answers during the action estimation task. Physical requests were instead implemented and controlled through the software environment RT-Lab, integrated with MATLAB/Simulink. RT-Lab included a 100 Hz loop for data storage, which permitted to collect the hand trajectories during the action execution task. The kinematic data recorded were analyzed using MATLAB (R2020b). Particularly, action velocity was estimated by using a third order Savitzky– Golay smoothing filter. Action velocity curves were obtained considering the time interval between the end of each request (starting position, Figure 1 B3.1) and the moment in which participants completed their passing action (ending position, Figure 1 B3.3).

### 2.3 Physical and vocal stimuli

As described in the previous section, participants could be physically or vocally stimulated before performing action duration estimation or action execution tasks. The physical parameters (velocity and trajectory) used to implement the physical request were derived from a previous kinematic recording in which an actor was asked to move the handle in a rude or gentle way. This procedure allowed us to generate a robotic movement that faithfully reproduced human vitality forms (rude and gentle). The stimuli performed through the manipulandum consisted of displacements on the horizontal plane starting from the coordinate (0m, -0.1m) of the workspace. For each physical request, the arm of participants was moved in the right direction and returned to the starting position. This robotic movement was completely different in terms of direction from those observed/executed by participants, avoiding a possible motor imitation effect between physical requests (frontal direction) and participants’ movements (rightward). Rude and gentle requests were presented in a random order and differed for trajectory and velocity. Additionally, in order to exclude a phenomenon of adaptation, for each vitality (rude and gentle) the manipulandum performed three movements with the same velocity (Figure 3 B2), but with a small angular shift among them (−10°, 0°, +10°; Figure 3 B1).

**Figure 3.**
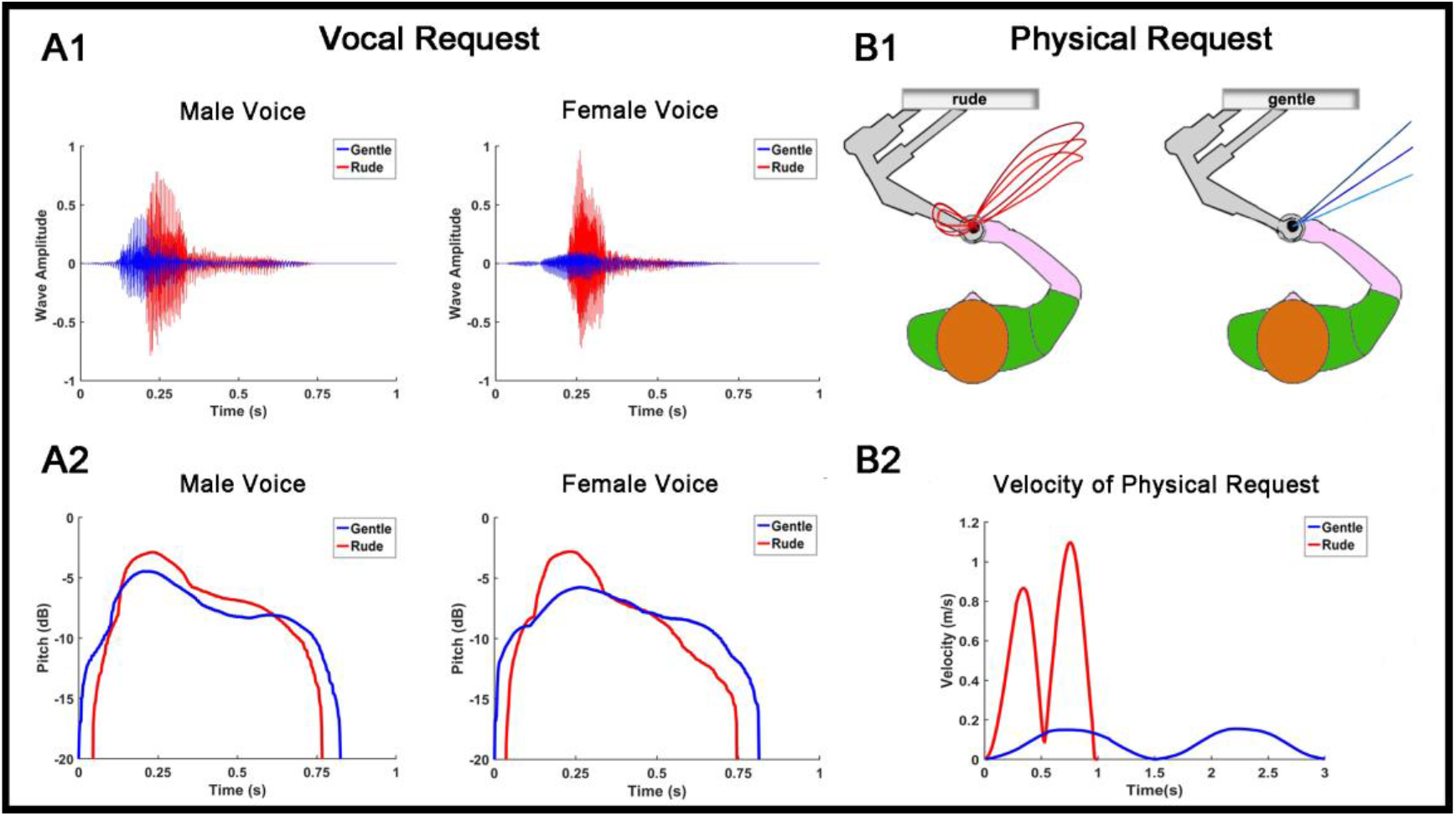
Wave amplitude (A1) and pitch (A2) of rude (red) and gentle (blue) vocal requests for the male and the female voice. Spatial trajectories of the motions performed by the manipulandum to provide rude (red lines) or gentle (blue lines) physical requests (B1). Velocity module of rude (red line) and gentle (blue line) physical requests (B2).

The rude request lasted 800ms with a maximum displacement of 22cm in the x-direction (right side). In contrast, the gentle request lasted 3000ms with a maximum displacement of 10cm in the same x-direction. On the other hand, during the vocal request participants listened to a male or a female voice pronouncing the Italian verb “dammi” (“give me”) in either a rude or a gentle way. Each vocal request was recorded using a condenser microphone (RODE NT1) placed 30 cm in front of the actors and digitized with a phantom powered A/D converter module (M-AUDIO M-TRACK). After recording, the audio files were processed with COOL EDIT PRO software. Rude and gentle vocal requests differed for parameters such as the wave amplitude and the pitch (Figure 3 A1, A2).

## 3. Results

### 3.1 Action Estimation Task

Considering the participants’ responses (estimated action duration) we measured the presence of possible differences between rude and gentle requests for both physical and vocal modalities. Specifically, the participants’ responses obtained after physical and vocal requests were normalized to the baseline condition (Estimated action duration(%) = estimated action duration after request *100 / estimated action duration during baseline condition), obtaining percentage values as shown in Figure 4. Then, four paired sample t-tests (two for gentle action estimation and two for rude action estimation) were carried out to assess possible differences between congruent and incongruent conditions, after physical (PHY) or vocal (VOC) requests. The significance level was fixed at p = 0.05. Before performing statistical analysis, the sphericity of data was verified (Mauchly’s test, p > 0.05). All variables were normally distributed (Kolmogorov-Smirnov Test, p > 0.05). Results showed a significant difference between congruent and incongruent conditions, for both gentle and rude vitality forms, regardless of the type of request (p < 0.05, for details see Figure 4AB).

**Figure 4.**
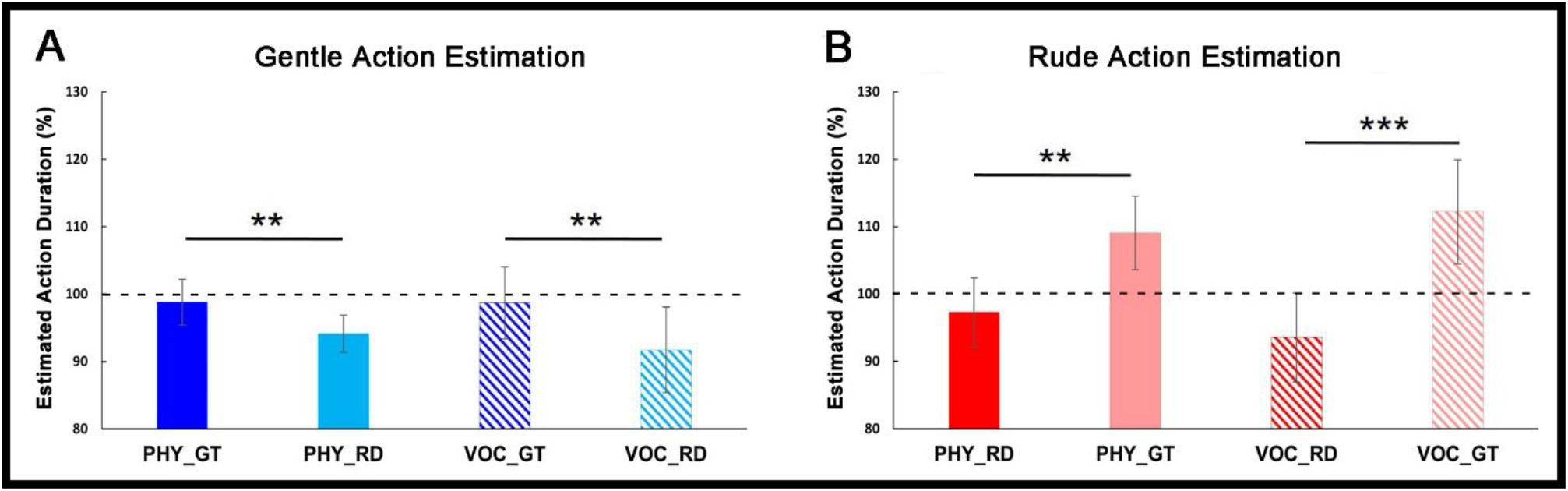
Results of gentle (A) and rude (B) action estimation. Estimated action durations normalized to the baseline condition are reported on the y-coordinate. The dotted line in correspondence of 100% refers to the baseline. PHY: physical request; VOC: vocal request; GT: gentle vitality form; RD: rude vitality form. Vertical bars represent the standard errors (SE). Horizontal bars indicate statistical significance (**p < 0.01, ***p < 0.001). Effect of rude requests on gentle action estimation

### 3.2 Action Execution Task

The action parameters characterizing the passage performed by participants after physical and vocal requests were normalized to the baseline condition as described above, obtaining percentage values as shown in Figure 5AB. Then, four paired sample t-tests (two for action velocity peak and two for distance covered after requests) were carried out to assess possible differences between actions performed by participants after physical (PHY) or vocal (VOC) requests. Results showed a significant difference between actions performed after rude and gentle requests, regardless the type of request (p <0.001 as shown in Figure 5AB). This difference is also highlighted in Figure 5CD, which shows the mean action velocity curves of participants in response to gentle and rude requests (physical and vocal). Furthermore, we quantify the effect of physical and vocal requests on participants’ motor responses regardless the vitality form conveyed. For this purpose, we calculated the effect of each request with respect to the baseline [PHY_RD – PHY_Baseline (100%)] and then mediated values obtained for the physical and the vocal request to obtain the overall effect of each modality. A paired sample t-test revealed a significant difference between physical and vocal requests (p<0.001). In particular, as shown in Figure 5EF, the effect of physical requests on action kinematic parameters is significantly higher than the effect of vocal requests (Velocity peak: PHY=26.5%, VOC=16.8%; Distance covered: PHY=18.6%, VOC=12.2%).

**Figure 5.**
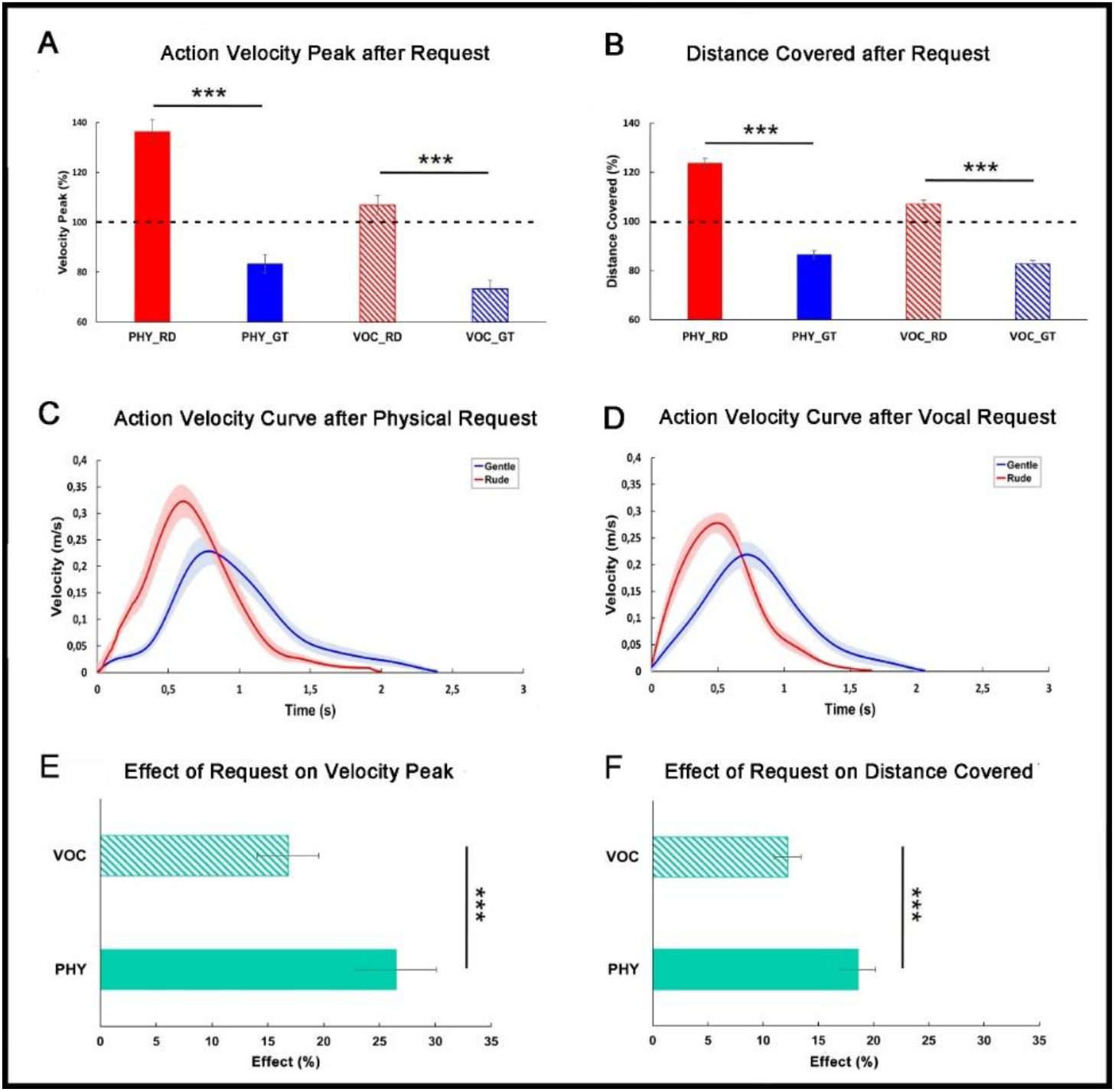
Action velocity peak (A) and distance covered (B) after request normalized to the baseline condition. Red color refers to rude requests (RD) and blue color refers to gentle requests (GT), both expressed physically (PHY) or vocally (VOC). Graphs in the middle show the action velocity curve characterizing the passage performed by participants after a physical (C) or a vocal (D) request. Error shading indicate standard error of the mean. At the bottom plots display the effect of physical (PHY) and vocal (VOC) request on modulating action velocity peak (E) and distance covered (F).

## 4. Discussion

Vitality forms represent a fundamental aspect of social communication allowing people to express their mood/attitude and to understand immediately those of others. For example, according to a positive or negative mood of the agent, an action can be unconsciously performed as gentle or rude towards another individual. In spite of their pervasiveness in human interactions, the influence of vitality forms on action perception has never been addressed. In this view, the aim of the present study is twofold: 1) to investigate whether and how vitality forms expressed physically or vocally by an agent may affect the participants’ responses during a cognitive task (action estimation task); 2) to assess how the same requests may modulate the kinematic parameters of a subsequent action (passing an object; action execution task). Results indicated that, during the action estimation task, a gentle request, independently of its modality (through physical contact or vocally) increased the duration of the subsequent action observed by participants. In contrast, a rude request affected the perception of an action subsequently presented, decreasing its perceived duration. More specifically, when participants observed the initial part of a gentle action but were previously stimulated with a rude request (incongruent condition), they anticipated the end of the action, even if it had a longer duration. On the other hand, if they observed the initial part of a rude action but they were previously stimulated with a gentle request (incongruent condition), they perceived the action as lasting longer, even if it had a shorter duration. Additionally, results of the action execution task indicated that, for both physical and vocal requests, the perception of vitality forms modulated the kinematic parameters (i.e., velocity and distance covered) of the subsequent action performed by participants. In particular, after a rude request, the action performed by participants (passing the object) had a higher velocity peak and covered a bigger distance. In contrast, after a gentle request, the action was performed with a lower velocity peak and covered a smaller distance. Notably, the effect produced by vitality forms during the action execution task was significant greater when the request was conveyed physically than vocally (see Figure 5CD). Our findings highlight the important role of vitality forms conveyed by action and speech in the modulation of the response to a social request, independently from the task and other social cues (e.g., facial expression). Interestingly, this modulation effect played by vitality forms on the receiver also occurred when participants listened to the vocal request, suggesting that the influence of vitality forms on the participants’ cognitive and motor tasks cannot be merely ascribed to a mechanism such as motor imitation. While several studies have shown the ability in encoding the goal of actions, our study represents the first attempt to demonstrate the ability in encoding the form of actions. Specifically, our results indicate that observing a very small part of an action, the observer is able to capture immediately the vitality form of that action. It is plausible that, during the tasks, participants processed the action kinematic information and remapped them on their own motor schema. This remapping process have probably permitted to represent internally the vitality forms of the observed action, preparing an adequate response.

A fundamental point of the present study is that, during social interactions, vitality forms expressed by the agent influence the behavior of the receiver. These findings are in agreement with another kinematic study, in which participants were presented with video clips showing two actors (1 actor and 1 actress) making gestural or verbal requests to pass a bottle (e.g., “give me”; giving task) or to act on it (“take the bottle”; taking task).^16^ Each request was presented as visual action, speech request, or both together (visual and speech) and was expressed gently and rudely. After the actors’ requests, participants performed the task (grasping a bottle with the goal to give or take it). Results showed that for both tasks (“giving” and “taking”), the speech and action vitality forms expressed by the actor’s vitality modulated the temporal (acceleration and velocity) and spatial parameters (trajectory) of a subsequent action performed by a receiver. In particular, concerning the reaching phase, vitality forms modulated acceleration, velocity, and trajectory of the reach component, showing a wider trajectory and higher velocity in response to the rude requests than to the gentle ones. Additionally, concerning the grasping phase, the results showed a wider maximal finger aperture in response to the rude vitality form than the gentle one. De Stefani and colleagues^19^ showed an analogous effect of prosody on the motor response of participants. Particularly, listening to the voice of an actress pronouncing positive or negative sentences, affected the kinematic parameters of a feeding motor action performed towards her. In line with our results, the affective component of vocal stimuli modulated the velocity of subsequent actions performed by participants, according to the negative or positive valence of words. Moreover, in a recent behavioral study Di Cesare and colleagues carried out two experiments in which participants listened to gentle/rude vocal requests and then observed the initial part of a passing action. Once the action was obscured, participants were required to continue it mentally, indicating its end (action estimation task of the present study).^20^ Differently from the present experiment, the authors used visual stimuli of different durations (200ms, 250ms, 300ms, 350ms for rude actions; 340ms, 420ms, 500ms, 600ms for gentle actions). Results showed that listening to rude/gentle vocal requests influenced the perception of actions subsequently observed. In addition, the same authors quantified the duration of this effect adding time delays (0ms, 200ms, 400ms, 800ms, 1200ms, 1600ms) between the vocal request and the video’s presentation, finding that the effect lasted 800ms and then started to decay.

Several fMRI studies have recently allowed highlighting neural correlates involved in vitality forms processing. More specifically, Di Cesare et al. showed that the perception (observation of actions, listening to action verbs) and expression (action execution, speech pronunciation) of different vitality forms produced the activation of the dorso-central insula. ^2-8^ These findings strongly suggest the existence of a mirror mechanism for action vitality forms in the dorso-central insula. Unlike the mirror mechanism located in the parieto-frontal circuit,^11,21-28^ which plays a role in action goal understanding, the mirror mechanism located in the dorso-central insula allows us to express our own vitality forms and to understand those expressed by others. Recently, Rizzolatti and colleagues showed that, besides actions and words, vitality forms could be also conveyed by tactile information. A typical example is handshaking that is a common greeting behavior in Western cultures. Authors showed that handshakes performed in an aggressive or gentle way produced, relative to a neutral control, stronger activation of the dorso-central insula.^17^ All together, these results suggest that the middle sector of the left insula is the key node selective involved in the processing of action, speech and tactile vitality forms. Concerning the present study, a possible interpretation is that the insula of the receiver encodes automatically the vitality forms of the request, preparing the appropriate response.

In some pathologies such as the autism spectrum disorder (ASD), children show impairment in vitality forms understanding.^29,30^ In a behavioral study, Rochat and colleagues^30^ asked children with ASD, as well as Typical Development children (TD), to observe video clips of actors (a male and a female) performing transitive and intransitive actions (e.g., passing a bottle and giving a high-five) in rude and gentle ways. Videos were presented in pairs, some pairs differing in the type of action (what task) others in the vitality forms (how task), and participants were required to judge whether video clips differed or not. Results showed that children with ASD, compared to TD, differed in the how task, while no difference was found in the what task. Recent data showed that although ASD children perform goal-directed actions correctly, they perform them in ways (i.e., with vitality forms) that are different from those of typically developing children.^31^ Taken together, these findings suggest that the perception and expression of vitality forms are essential skills that we need to acquire in order to communicate with others.

Interestingly vitality forms can be effective also when expressed by non-human agents. Drawing inspiration from human voice and movement, a recent study^32^ modified a passing action and a voice generated by the humanoid robot iCub, with the aim of transmitting vitality forms. Results showed that the kinematic parameters of the robot’s movement and the properties of its voice are adequate to express different attitudes, which are consistently perceived as rude or gentle by human partners. Moreover, the same authors showed that the observation of robotic actions conveying vitality forms (rude and gentle) produced an increase of the BOLD signal in the dorso-central insula, the same sector activated during the perception of vitality forms expressed by humans.^33^

In conclusion, our data shed new light on the fundamental role of vitality forms in social interactions. We clearly demonstrated how physical and vocal requests conveying different vitality forms modulate the response of the receiver. In particular, when an individual asks us something, his/her positive and negative attitudes, communicated by vitality forms, modulate both our perception and motor behavior. The role of vitality forms in influencing others from an affective state point of view highlights their relevance in social communication. Results and methodology from this study may have implications for social and communicative disorders and other research fields such as robotics.

## Acknowledgements

This work has been supported by a Starting Grant from the European Research Council (ERC) under the European Union’s Horizon 2020 research and innovation programme. G.A. No 804388, wHiSPER

## Author Contributions

GDC, JZ, AS and FR designed the research; GDC and GL performed the experiment and analyzed the data; JZ, FV and GB contributed with material and consulting for the development of the project; GDC and GL wrote the first draft, all authors read the manuscript and contributed to its final form.

